# Trehalose exerts cryoprotective effects on piglet testicular tissue

**DOI:** 10.64898/2026.08.18.745558

**Authors:** Yuanyuan Huang, Nian Liu, Juan Liu, Yixian Wei, Xiaoye Wang, Xun Li, Jiaming Zheng, Changlong Xu, Chuanhuo Hu

## Abstract

The cryopreservation of testicular tissue is crucial for maintaining male fertility; However, its efficacy is often compromised by oxidative stress and mitochondrial dysfunction. Trehalose, a natural cryoprotectant, demonstrates significant potential, yet its specific mechanisms, particularly in mitochondrial regulation, remain insufficiently characterized. This study aimed to investigate the cryoprotective effects of trehalose on testicular tissue from 18-21-day-old piglets, with a focus on mitochondrial metabolism. Samples were cryopreserved via a slow-freezing protocol in a modified standard solution containing 200 mmol/L trehalose. The protective effect was evaluated by measuring testosterone synthesis, blood testis barrier (BTB) and spermatogenesis. Additionally, protective outcomes were assessed by measuring cell viability, tissue morphology, reactive oxygen species (ROS) levels, apoptosis rates, and testosterone secretion following freeze-thaw cycles. Transcriptomic sequencing and bioinformatics analyses were conducted to elucidate the underlying molecular mechanism. Cryopreservation led to reduced testosterone synthesis and secretion, decreased levels of BTB-binding proteins, and impaired spermatogenesis. Results indicated that 200 mmol/L trehalose significantly improved cell survival, decreased apoptosis and ROS levels, and enhanced testosterone secretion. 200 mmol/L trehalose partially increased the expression of *StAR* and *CYP11A1* genes associated with testosterone synthesis while it protected the tight junction proteins Claudin-11, ZO-1 and the gap junction protein Cx43. Consequently, it exerted a reproductive protective effect by increasing the expression of key spermatogenic regulators *DDX25*, *HMGB2*, acrosomal protein *DYP19L2*, and sperm tail proteins AKAP4 and CFAP44. Transcriptomic profiling demonstrated that trehalose predominantly restored the transcriptional expression of genes involved in the mitochondrial electron transport chain and oxidative phosphorylation pathways, including ND2, COX2, ATP8, ATP6, ND5, ND6 and CYTB. These findings indicate that trehalose primarily protects piglet testicular tissue during cryopreservation by enhancing mitochondrial function, thereby providing a molecular basis for optimizing cryopreservation protocols.

## 1. Introduction

Testicular tissue cryopreservation is a vital strategy for preserving male genetic resources, particularly in prepubertal animals or those that die unexpectedly before sperm production begins [1]. Unlike sperm cryopreservation, which is limited to sexually mature males, testicular tissue cryopreservation allows for the preservation of spermatogonial stem cells (SSCs) and the testicular microenvironment, enabling future applications such as transplantation, in vitro culture, and assisted reproductive technologies [2]. However, cryopreservation can cause severe damage to testicular tissue, including ice crystal formation, oxidative stress, and mitochondrial dysfunction, leading to reduced cell viability and impaired tissue function [3,4].

Trehalose, a naturally occurring disaccharide, has emerged as an effective cryoprotectant due to its unique properties, including high hydrophilicity, chemical stability, and antioxidative activity. It has been extensively used for cryopreservation of sperm, SSCs, and testicular tissue across various species [5–7]. Lee et al. [8] found that 200 mmol/L trehalose to serum-free freezing medium effectively preserves mouse SSCs with their proliferative capacity and stem cell activity following transplantation maintained. Trehalose to diluted bovine semen alleviated oxidative stress damage caused by the freezing and thawing process and improves semen quality [9]. Trehalose protected cells and tissues by stabilizing membranes and proteins, reducing oxidative stress, and maintaining mitochondrial function [10]. Pigs, due to their physiological similarities to humans, serve as important agricultural and biomedical models. However, the effects of trehalose on pre-pubertal piglet testicular tissue cryopreservation, particularly on mitochondrial function, remain poorly understood.

Mitochondria play a crucial role in cellular energy production, steroidogenesis, and apoptosis regulation. Freezing-induced mitochondrial dysfunction may lead to excessive reactive oxygen species (ROS) production, oxidative damage, and impaired testosterone synthesis [2,11,12]. Mitochondrial activity and mitochondrial ROS decreased during the cryopreservation of boar semen; this was attributed to an increase in soluble mitofusin-2 and a decrease in actin, which led to abnormal mitochondrial volume [13]. Xi et al. [14] found that treatment with trehalose increases the expression of the mitochondrial marker COX IV with the expression of the fusion protein MFN1, the fission protein p-Drp1/Drp1, and the mitochondrial biogenesis proteins TOM20 and PGC1-α upregulated, thereby improving mitochondrial dysfunction in mice with D-galactose-induced testicular aging. Therefore, protecting mitochondrial function during cryopreservation is essential for maintaining the viability and functionality of testicular tissue.

In this study, 200 mmol/L trehalose was used to investigate its effects on testosterone synthesis and secretion, BTB integrity, and gene expression related to spermatogenesis. Subsequently, transcriptomic profiling were employed to identify key genes and pathways regulated by trehalose, with the aim of validating the hypothesis that trehalose exerts its cryoprotective effects by enhancing mitochondrial function.

## 2. Materials and Methods

### 2.1. Testicular Tissue Collection and Cryopreservation

Testes were collected from 30 pre-pubertal male piglets (18–21 days old) at a commercial farm in the suburban area of Nanning, China. Following surgical castration under intravenous anesthesia, postoperative care included suturing, antisepsis, and antibiotic prophylaxis. Piglets recovered fully and were returned to standard herd management until market weight. Excised tissues were transported to Guangxi University for analysis. All animal housing and experimental procedures were strictly conducted in accordance with the guidelines established by the Institutional Animal Care and Use Committee (IACUC) of Guangxi University (Approval No. Gxu2022-211). After washing, the tissue was cut into 2-3 mm³ fragments. The base freezing medium consisted of DMEM (11965092, Gibco, Grand Island, NY, USA) supplemented with 10% DMSO (ST038, Beyotime, Shanghai, China), 10% fetal bovine serum (FBS, 10437028; Gibco, Grand Island, NY, USA), and 5% penicillin-streptomycin (C0222; Beyotime, Shanghai, China). To determine the optimal concentration of trehalose (T9531, Sigma, St. Louis, MO, USA), testicular tissues were cryopreserved in media supplemented with graded concentrations of trehalose (50, 100, 200, 400, and 800 mmol/L). A Fresh group (non-frozen) and a cryoprotectant-only Control group (0 mmol/L trehalose) were established as references. Following a 1 h equilibration in freezing medium at 4°C, tissue fragments were transferred to -20°C for 2 h, then to -80 ° C for overnight storage. After a minimum of 7 days in liquid nitrogen, samples were retrieved for analysis. The Fresh group samples were maintained at 4°C without freezing.

### 2.2. Preparation of Single-cell Suspensions and Cell Viability Assessment

Testicular single-cell suspensions were prepared under sterile conditions: thawed tissues were washed thrice with 10× PBS (1500 rpm, 5 min) until supernatants cleared, sequentially digested with Tissues were sequentially digested with collagenase IV (C5138, Sigma, St. Louis, MO, USA) at 1 mg/mL for 30 min, followed by 0.25% trypsin (T8003, Sigma, St. Louis, MO, USA) for 20 min at 37°C, and terminated with equal-volume DMEM containing 10% FBS. Filtrates through a 40 μm strainer were centrifuged (2000 rpm, 5 min), washed twice, and resuspended in complete DMEM. Cell viability was assessed by trypan blue exclusion: suspensions adjusted to 1 × 10⁶ cells/mL were mixed with equal-volume 0.4% trypan blue (C1313L, Beyotime, Shanghai, China), counted within 3 min via hemocytometer (≥ 500 cells per sample), and viability calculated as (viable cells/total cells)× 100%.

### 2.3. Testicular Cell Morphology Analysis

For histological evaluation, testicular tissue fragments from all groups were fixed in 4% paraformaldehyde, routinely processed for paraffin embedding, and sectioned at 5 μm. Sections were stained with hematoxylin and eosin (H&E) and examined under a bright-field microscope (Eclipse E100; Nikon, Tokyo, Japan). Morphological parameters including tubular structure, basement membrane integrity, germ cell swelling, cell loss, basement membrane detachment, and vacuolization were scored on a 0–3 scale according to Sliva et al. [15] (0 = severe damage, 3 = normal morphology; Table 2-1). At least 30 seminiferous tubules were randomly evaluated per group in a blinded manner.

**Table 2-1.** Morphological parameters of seminiferous tubules from the porcine testes.

| Parameter | Score |  |  |
| --- | --- | --- | --- |
|  | #3 | #2 | #1 |
| Tubular cell swelling | No swelling | > 50% cells without swelling | > 50% cells with swelling |
| Tubular cell loss | No cell loss | < 75% cell types lost | > 75% cell types lost |
| Rupture from basal membrane | No rupture | Partly rupture (< 50%) | Mostly Rupture (> 50%) |

### 2.4. TUNEL Assay for Apoptosis Detection

Apoptosis was detected using the In Situ Cell Death Detection Kit, Fluorescein (#11684795910, Roche Diagnostics, Mannheim, Germany) according to the manufacturer’s protocol. Following deparaffinization and rehydration, sections were permeabilized with 20 μg/mL proteinase K (ST532, Beyotime, Shanghai, China) for 15 min, after which endogenous peroxidase activity was quenched with 0.3% hydrogen peroxide in methanol for 15 min. Sections were then blocked with 5% bovine serum albumin (BSA; 4240GR100, BioFroxx, Einhausen, Germany) for 1 h at room temperature, prior to incubation with 50 μ L of the TUNEL reaction mixture at 37 ° C for 1 h in the dark. Nuclei were counterstained with DAPI (C1006, Beyotime, Shanghai, China) for 10 min, and sections were mounted with an anti-fade mounting medium (AR0036, Boster, Wuhan, China). Fluorescence imaging was performed on a BX53 fluorescence microscope (Olympus, Tokyo, Japan) using 450–500 nm excitation and 515–565 nm emission filters. For each sample, ten random non-overlapping fields were acquired at 400× magnification, with at least one seminiferous tubule per field. Apoptosis rates were calculated as the percentage of TUNEL-positive cells relative to total DAPI-positive nuclei, with 300–600 cells counted per sample.

### 2.5. ROS Assay

Intracellular ROS was measured using a ROS Assay Kit (S0033S, Beyotime, Shanghai, China). Cell pellets were resuspended in 1:1000 DCFH-DA/PBS, incubated at 37°C in the dark for 30 min with 5-min inversion. After three PBS washes (1500 rpm, 5 min), cells were resuspended in 200 uL PBS. Images were captured at 200× under a fluorescence microscope (Olympus, Tokyo, Japan); ten random fields per sample were analyzed via ImageJ, with ROS levels normalized to the Fresh group.

### 2.6. In Vitro Culture of Testicular Tissue and Measurement of Testosterone Secretion

Fresh or thawed testicular tissue was washed with PBS and placed in a culture dish. The base freezing medium was added to ensure complete coverage of the tissue. The dish was incubated in an incubator at 37°C and 5%CO₂ for 24 h. The medium was collected, and testosterone secretion levels were measured using a sandwich immunoassay kit (JM-061493O1, Jingmei biotechnology, Shenzhen, China).

### 2.7. RNA Extraction and qRT-PCR Assay

For validation assays, total RNA was extracted from testicular tissues using RNA Isolater (R401-01, Vazyme, Nanjing, China) according to the manufacturer’s protocol for animal tissues. cDNA was synthesized using the HiScript II 1st Strand cDNA Synthesis Kit (R211, Vazyme, Nanjing, China) as per the manufacturer’s instructions, with incubation at 25°C for 5 min, 50°C for 15 min, and 85°C for 2 min. The resulting cDNA was either immediately used for qPCR or stored at -20°C.

qPCR Reactions were performed using the 2× SYBR Green qPCR Mix (AH0101, Sparkjade, Jinan, China). The 20 μL reaction mixture comprised: 10 μL 2× SYBR Green qPCR Mix, 1 μL cDNA template, 0.5 μL forward primer (10 μM), 0.5 μL reverse primer (10 μM), and 8 μL RNase-free water. The thermal cycling protocol consisted of an initial denaturation at 94°C for 3 min, followed by 40 cycles of 94°C for 10 s and 60°C for 30 s. Based on the existing literature, genes that are widely recognized as key markers for testosterone synthesis, BTB function, and spermatogenesis were selected: *StAR*, *CYP17A1*, *CYP11A1*, *Claudin-11*, *ZO-1*, *Cx43*, *AKAP4*, *CFAP44*, *DDX25*, *DPY19L2*, *HMGB2* and *GAPDH* [16–18]. Primers were designed using Primer Premier 5.0 software based on pig mRNA sequences available in the NCBI database and synthesized by Nanning Qingke Biotechnology Co., Ltd. Additionally, ten differentially expressed genes (*PSMC6*, *PSMA3*, *NFE2L2*, *ND2*, *COX2*, *ATP8*, *ATP6*, *ND5*, *ND6*, and *CYTB*) were selected to validate the RNA-seq results; their primer sequences are listed in Table S1. The relative expression level of the target gene was calculated using the 2^-△△ct^ method.

### 2.8. Protein Isolation and Western Blot

Briefly, frozen-thawed testicular tissues were weighed and homogenized in ice-cold RIPA (#9806, Cell Signaling Technology, Danvers, MA, USA) lysis buffer containing PMSF (#8553, Cell Signaling Technology, Danvers, MA, USA) protease inhibitor at a ratio of 100 mg tissue per 1 mL buffer. The homogenate was incubated at 4°C for 30 min and then centrifuged at 12,000 × g for 10 min at 4°C. The supernatant was collected, and the protein concentration was determined using a BCA (PC0021, Solarbio, Beijing, China) assay. Protein samples were denatured by boiling with loading buffer at 100°C for 10 min and stored at -20°C.

For Western blotting, proteins were separated using either a 12.5% SDS-PAGE Denaturing Acrylamide Gel Rapid Preparation Kit (C681103, Sangon Biotech, Shanghai, China) or an 8% ExpressCast PAGE Color Gel (P2011, NCM Biotech, Suzhou, China). Equal amounts of protein extract (25 µg per lane) were transferred onto a polyvinylidene fluoride (PVDF) membrane (IPVH00010, Merck Millipore Ltd, Cork, Ireland). Membranes were blocked for 2 h with 5% non-fat dry milk in PBST and incubated overnight at 4°C with primary antibodies against CX43 (rabbit, 1:5000, 80543-1, Proteintech, Rosemont, IL, USA), ZO-1 (rabbit, 1:1000, TB7663, Abmart, Shanghai, China), AKAP4 (rabbit, 1:1000, A14813, Abclonal, Wuhan, China), and HMGB2 (rabbit, 1:750, A17360, Abclonal, Wuhan, China). Subsequently, the membranes were incubated with appropriate HRP-conjugated secondary antibodies. Protein bands were visualized using an enhanced chemiluminescence (ECL, AR0091, Boster, Wuhan, China) substrate, and the band intensities were quantified with Image J software.

### 2.9. Quality Assessment of Transcriptome Sequencing Data

Transcriptome analysis to nine samples was underwent, producing 77.53 Gb of clean data in total. Each sample had more than 6.06 Gb of data after quality control, with a base error rate of less than 0.1%, a Q20 of more than 85%, and a Q30 base percentage of more than 80%. All samples maintained a GC content of almost 50%. This suggests that the sequencing data (Table S2) is of excellent quality and appropriate for further examination.

### 2.10. Differential Expression Analysis

Raw reads were processed to remove adapters and low-quality sequences. Clean reads were mapped to the reference genome using HISAT2. Gene expression levels were quantified using FPKM (Fragments Per Kilobase of transcript per Million mapped reads). Differentially expressed genes (DEGs) were identified using DESeq2 with a threshold of |log2FC| > 1.5 and *P* adjust < 0.05.

### 2.11. GO and KEGG Enrichment Analysis

Gene Ontology (GO) enrichment analysis was performed to identify biological processes (BP), cellular components (CC) and molecular functions (MF) associated with the DEGs. KEGG pathway analysis was conducted to explore the signaling pathways involved. Enrichment was considered significant at *P* adjust < 0.05.

### 2.12. Data Analysis

All experiments were conducted with at least three independent biological replicates per treatment group (n = 3). One-way analysis of variance (ANOVA) to all results was underwent using SPSS 24.0 statistical software, with graphical representations created using GraphPad Prism 8. Data were expressed as mean ± standard deviation. *P* < 0.05 indicated significant differences.

## 3. Results

### 3.1. The Effect of Trehalose on Post-Thaw Cell Viability

Total cell viability was determined by trypan blue exclusion (membrane integrity loss = cell death). All cryopreserved groups showed significantly lower viability than the Fresh group (*P* < 0.01, Figure. 3-1). Trehalose supplementation (50–800 mmol/L) significantly improved viability relative to the Control group (*P* < 0.01), with 200 mmol/L trehalose yielding peak viability across all treatment groups (*P* < 0.01).

**Figure 3-1.**
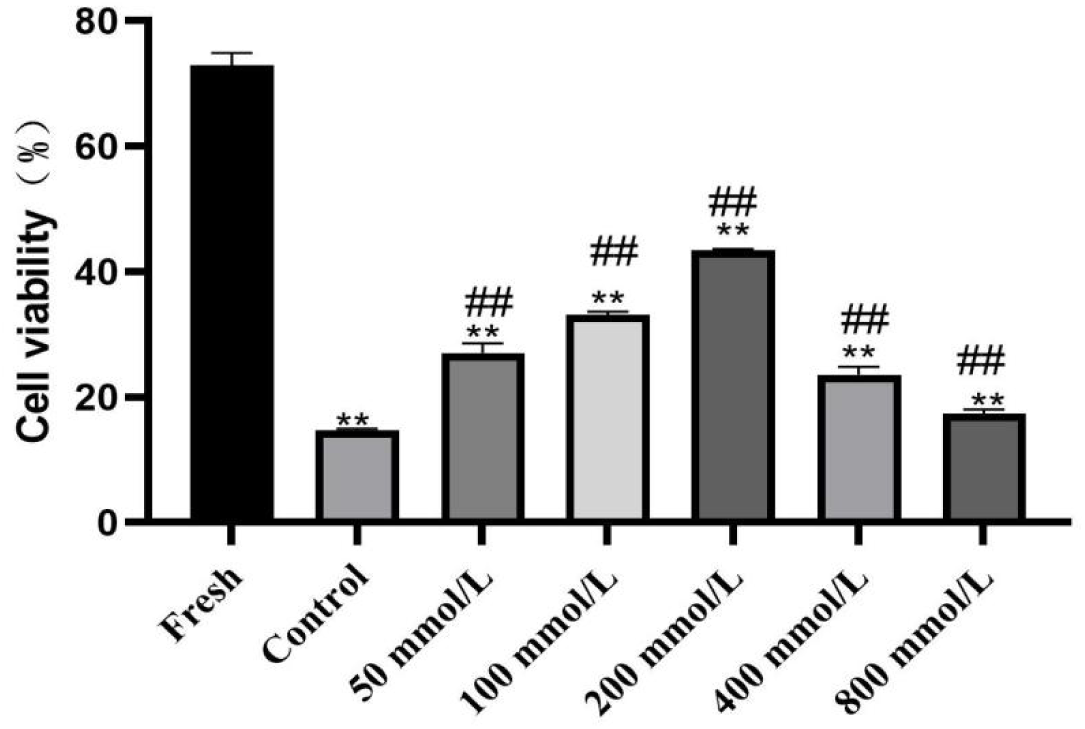
Total cell viability of the fresh and frozen-thawed piglet testicular tissue. The data are means ± SD (n = 3). \*\**P <* 0.01 indicates a statistically significant difference vs Control-800 mmol/L trehalose groups; ^##^*P <* 0.01 indicates a statistically significant difference vs 50-800 mmol/L trehalose groups.

### 3.2. The Effect of Trehalose on Post-Thaw Morphology of Piglet Testicular Tissues

The histomorphological features of fresh and frozen testicular tissues are presented in Figure. 3-2. Overall, in Figure. 3-3 the Fresh group and 200 mmol/L trehalose groups exhibited comparable morphological scores across all parameters (*p* > 0.05). Specifically, supplementation with 200 mmol/L trehalose better preserved seminiferous tubule integrity and prevented cellular swelling compared to the 50 mmol/L trehalose and the Control groups (*p* < 0.05). While the Fresh group maintained optimal tubular architecture, the 200 mmol/L trehalose group demonstrated significantly reduced germ cell loss and basement membrane rupture relative to the 100 mmol/L trehalose and the Control groups (*p* < 0.01). Conversely, lower (50 mmol/L trehalose) and higher (400/800 mmol/L trehalose) concentrations offered no significant protection against cryo-injury compared to the Control group (*p* > 0.05).

**Figure 3-2.**
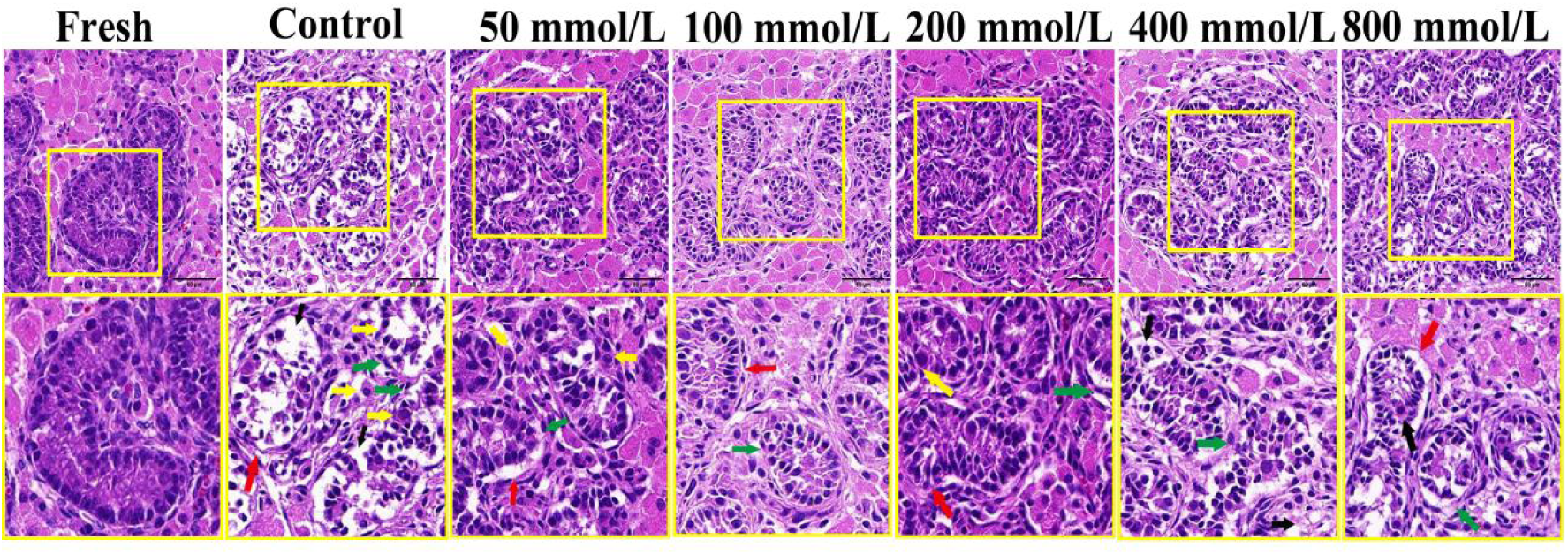
Histological images of the fresh and frozen-thawed piglet testicular tissue (×400, scale bar 50 µm). Yellow arrows indicate tubular cell swelling; Black arrows indicate tubular cell loss; Red arrows indicate shrinkage from basal membrane; Green arrows indicate rupture from basal membrane.

**Figure 3-3.**
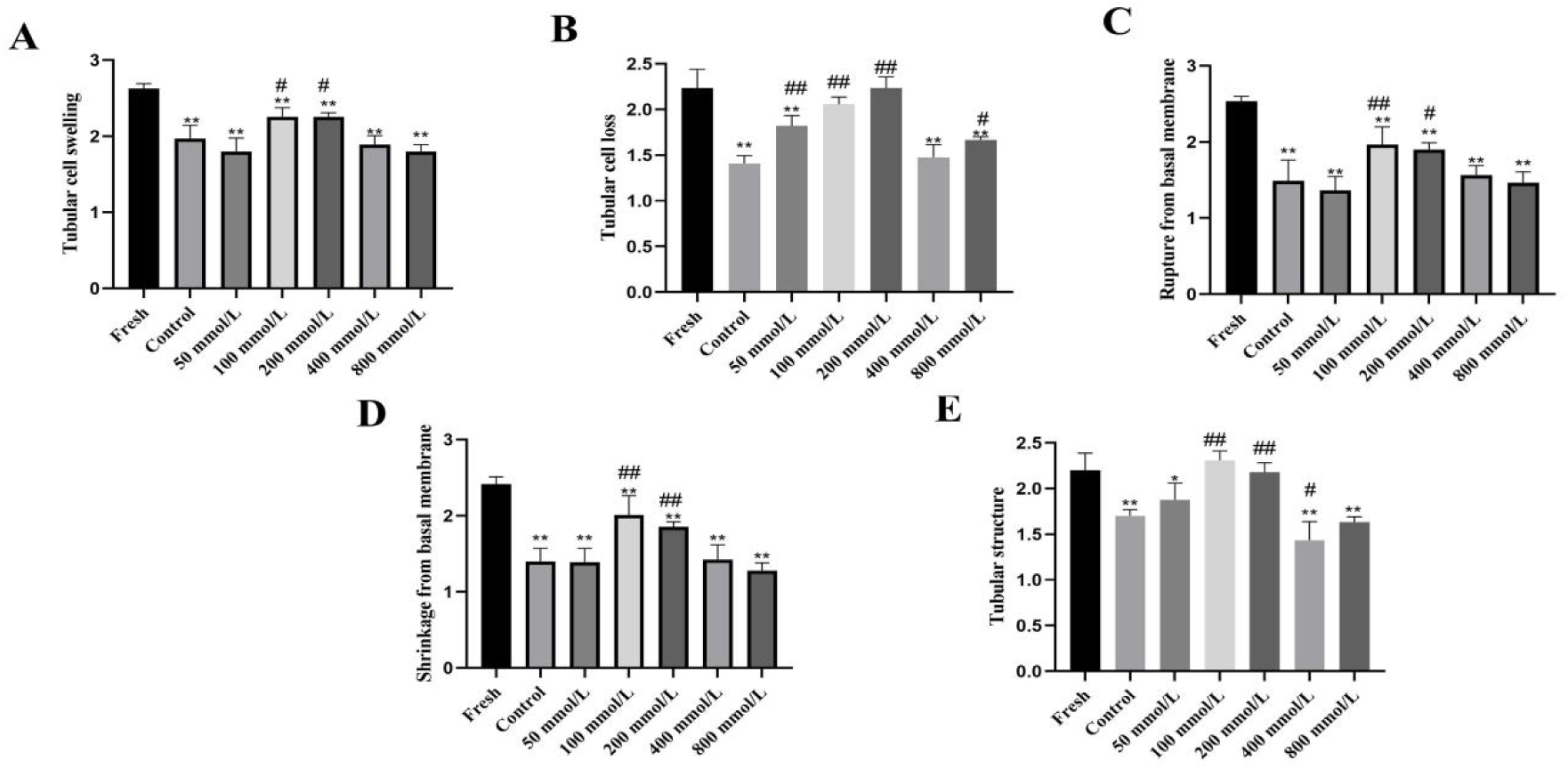
Morphological evaluation of the fresh and frozen-thawed piglet testicular tissue. (A) Tubular cell swelling; (B) Tubular cell loss; (C) Shrinkage from basal membrane; (D) Rupture from basal membrane; (E) Tubular structure. The data are means ± SD (n = 3). \**P <* 0.05 and \*\**P <* 0.01 indicates a statistically significant difference vs Control-800 mmol/L trehalose groups. ^#^*P <* 0.05 and ^##^*P <* 0.01 indicates a statistically significant difference vs 50-800 mmol/L trehalose groups.

### 3.3. The Effect of Trehalose on Apoptosis in Post-Thaw Porcine Testicular Tissue

Representative TUNEL staining images are shown in Figure. 3-4A. Overall, the apoptosis rate of the 200 mmol/L trehalose group was comparable to that of the Fresh group (*p* > 0.05), while the 100 mmol/L trehalose group showed a significant elevation versus the Fresh group (*p* < 0.05); all other trehalose-treated groups exhibited markedly higher apoptosis rates relative to the Fresh group (*p* < 0.01). Compared with the Control group, supplementation with 50, 100 and 200 mmol/L trehalose significantly reduced the apoptosis rate (*p* < 0.01), whereas higher concentrations (400 and 800 mmol/L trehalose) conferred no statistically significant protection (*p* > 0.05, Figure. 3-4B).

**Figure 3-4.**
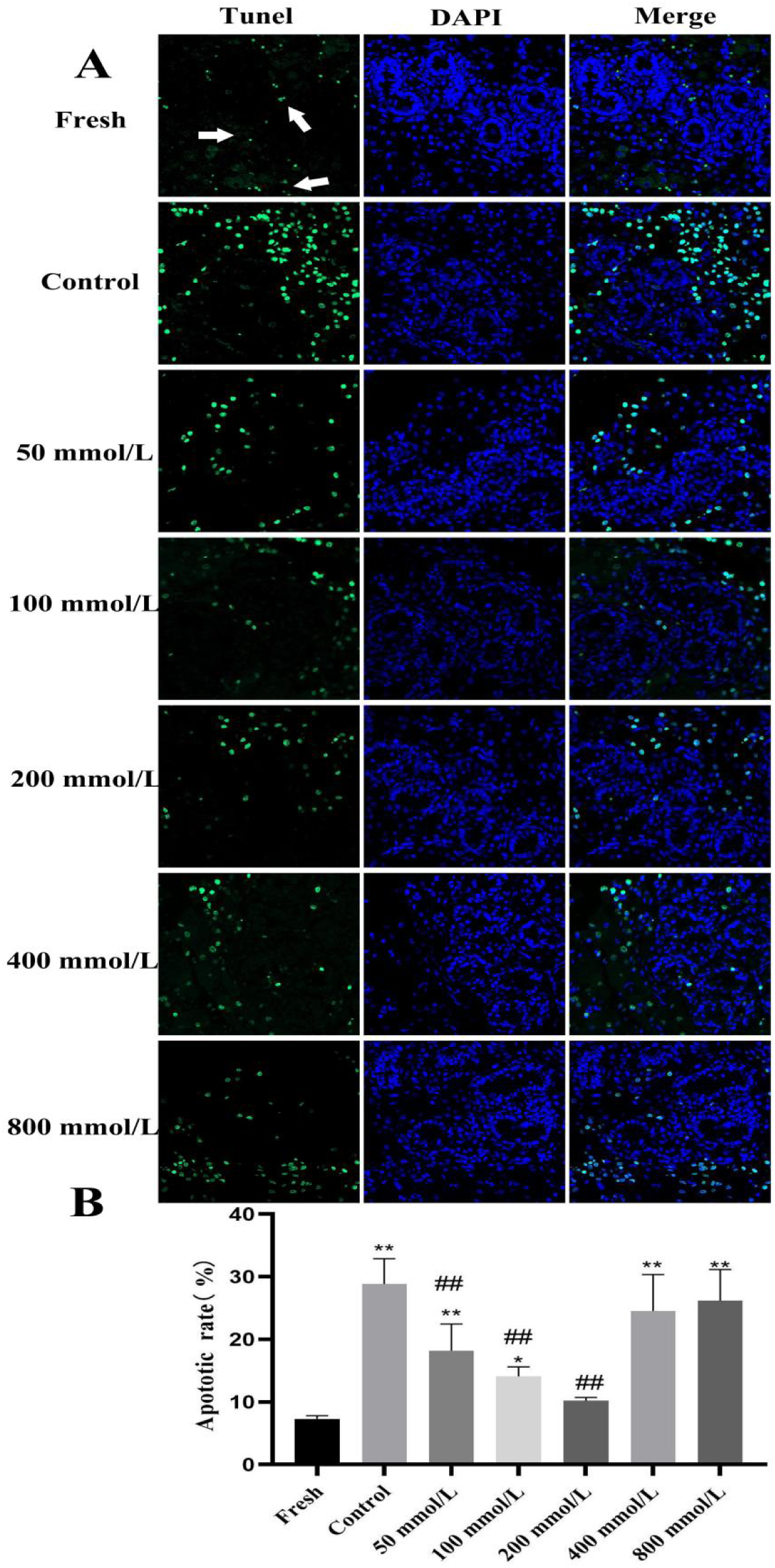
Apoptotic analysis of the fresh and frozen-thawed piglet testicular tissue (×400). White arrows indicate apoptotic cells. (A) Representative TUNEL staining of each group; (B) corresponding apoptosis rates. The data are means ± SD (n = 3). \**P* < 0.05 and \*\**P* < 0.01 indicates a statistically significant difference vs Control-800 mmol/L trehalose groups; ^##^*P* < 0.01 indicates a statistically significant difference vs 50-800 mmol/L trehalose groups.

### 3.4. The Effect of Trehalose on ROS Levels in Post-Thaw Porcine Testicular Tissue

Intracellular ROS levels were assessed using the DCFH-DA probe, which is oxidized by intracellular ROS to generate fluorescent DCF, with fluorescence intensity reflecting ROS abundance (Figure. 3-5A). Overall, all cryopreserved groups exhibited markedly elevated ROS levels compared to the Fresh group (*p* < 0.01). Specifically, supplementation with 50 mmol/L trehalose (*p* < 0.05), 100 mmol/L trehalose (*p* < 0.01), 200 mmol/L trehalose (*p* < 0.01) and 800 mmol/L trehalose (*p* < 0.05) trehalose significantly reduced ROS relative to the Control group, whereas the 400 mmol/L trehalose group showed no statistically significant difference versus the Control group (*p* > 0.05, Figure. 3-5B).

**Figure 3-5.**
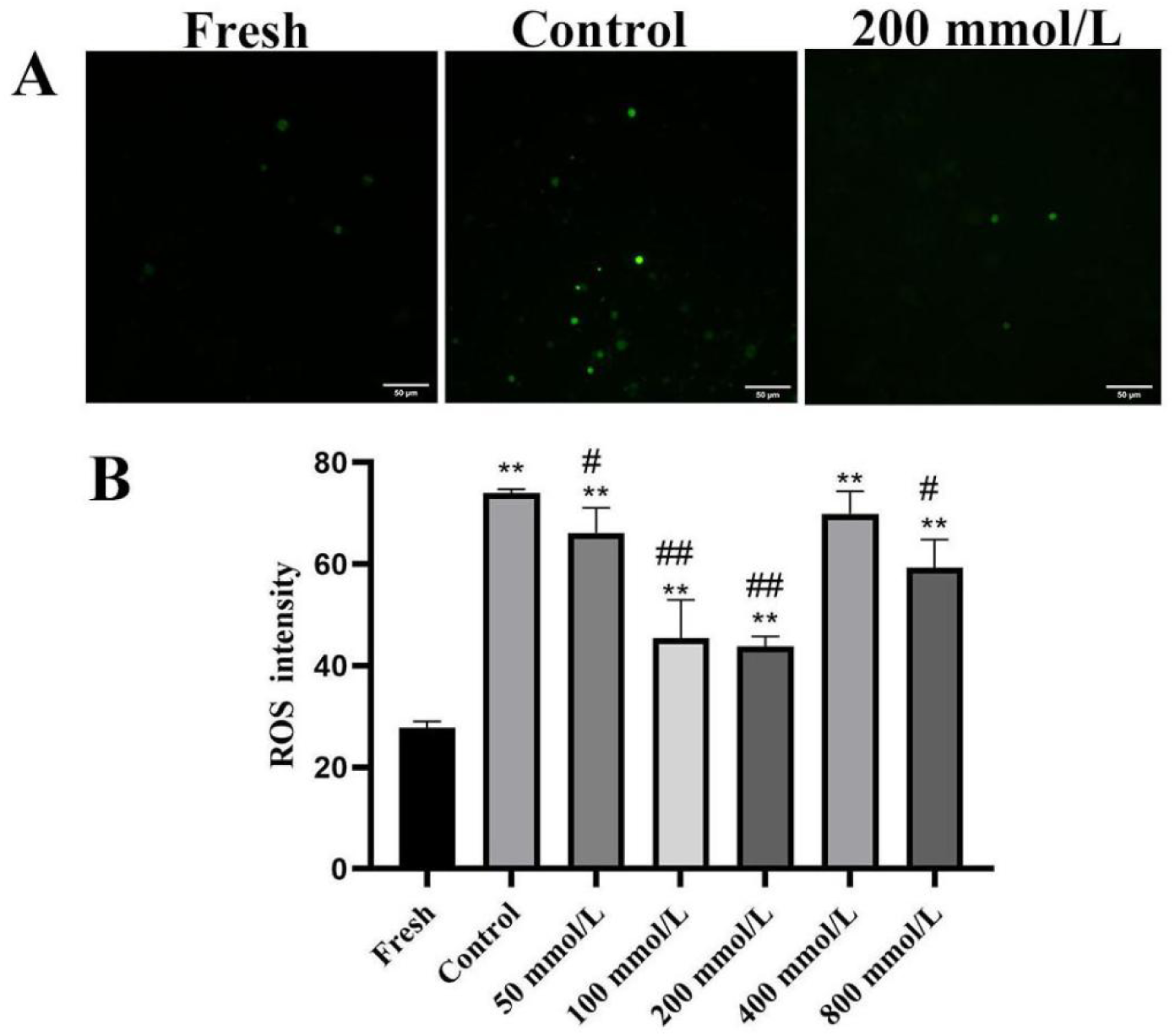
Statistical results of ROS of the fresh and frozen-thawed piglet testicular tissue. (A) indicate the fluorescence emitted by cells under a fluorescence microscope (×200, scale bar 50 µm); (B) indicate the fluorescence intensity of ROS at different concentrations of trehalose. The data are means ± SD (n = 3). \*\**P* < 0.01 indicates a statistically significant difference vs Control-800 mmol/L trehalose groups; ^#^*P* < 0.05 and ^##^*P* < 0.01 indicates a statistically significant difference vs 50-800 mmol/L trehalose groups.

### 3.5. The Effects of Trehalose on Testosterone Synthesis and Secretion in Piglet Testicular Tissue

Testosterone is one of the key hormones synthesized by Leydig cells (LCs). First, ELISA was used to measure testosterone secretion in the culture medium from frozen-thawed testicular tissue. The results are shown in Figure 3-6A. Compared with the Fresh group, testosterone secretion in Control group culture medium was significantly reduced (*P* < 0.01); Subsequently, qRT-PCR was used to detect the mRNA expression of *CYP17A1*, *CYP11A1*, and *StAR*, genes related to testosterone synthesis. As shown in Figure 3-6B-D, the mRNA expression of *CYP17A1*, *CYP11A1*, and *StAR* in frozen-thawed piglet testicular tissue all showed a significant decrease (*P* < 0.01); Compared with the Control group, although there was no significant difference in *CYP17A1* expression, 200 mmol/L trehalose significant increased the testosterone secretion (*P* < 0.05) and *CYP11A1* and *StAR* mRNA expression (*P* < 0.01).

**Figure 3-6.**
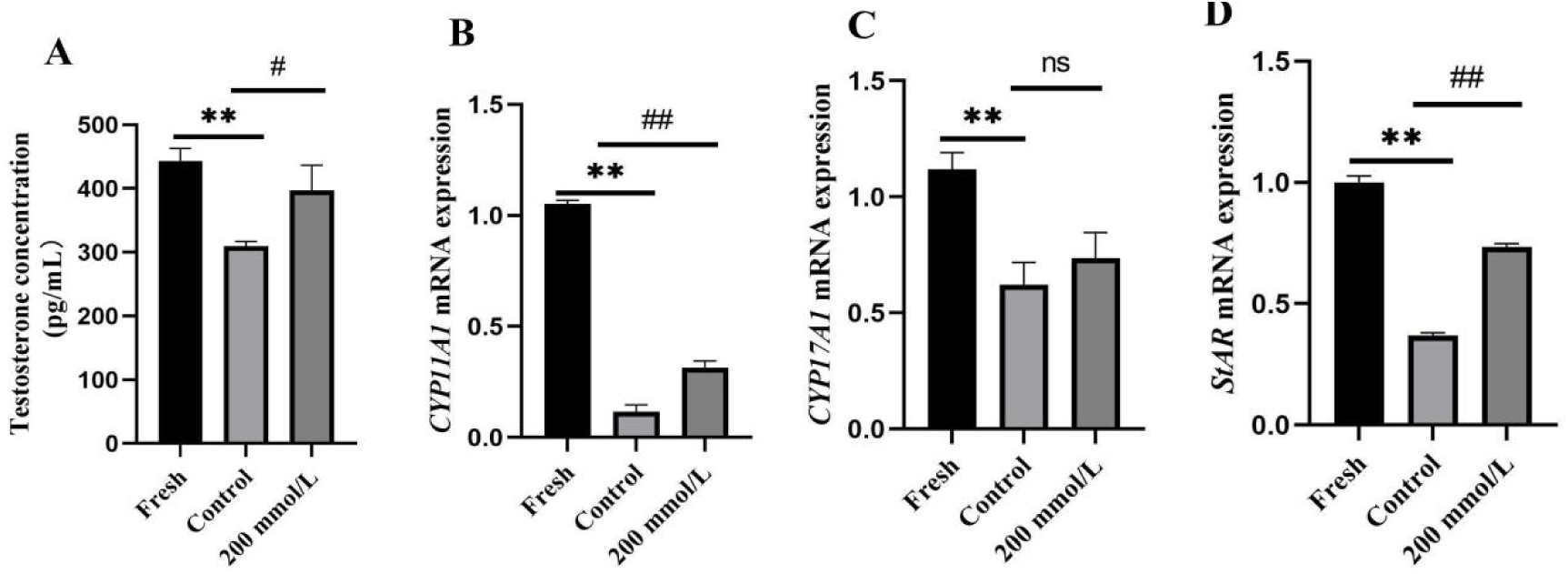
Effect of trehalose on testosterone synthesis of frozen-thawed piglet testicular tissue. (A) Testosterone concentration; (B) *CYP11A1* mRNA expression; (C) *CYP17A1* mRNA expression; (D) *StAR* mRNA expression. The data are means ± SD (n = 3). \*\**P* < 0.01 indicates a statistically significant difference vs Fresh group; ^#^*P* < 0.05 and ^##^*P* < 0.01 indicates a statistically significant difference vs Control group.

### 3.6. The Effects of Trehalose on BTB of frozen-thawed Piglet Testicular Tissue

Tight junctions formed by adjacent SCs are a major component of the BTB, which play a crucial role in maintaining the testicular microenvironment for spermatogenesis. qRT-PCR and Western blot were used to detect the mRNA expression of *Claudin-11*, *Cx43*, and *ZO-1*, as well as the protein expression of Cx43 and ZO-1, respectively. The results are shown in Figure 3-7. Compared with the Fresh group, the mRNA expression of *Claudin-11*, *Cx43*, and *ZO-1* in frozen-thawed piglet testicular tissue was significantly reduced (*P* < 0.01), and the protein expression of ZO-1and Cx43 was also significantly reduced (*P* < 0.01); Compared with the Control group, 200 mmol/L trehalose significantly increased the mRNA expression of *Claudin-11*, *Cx43* and *ZO-1*, and the protein expression of Cx43 and ZO-1 (*P* < 0.01).

**Figure 3-7.**
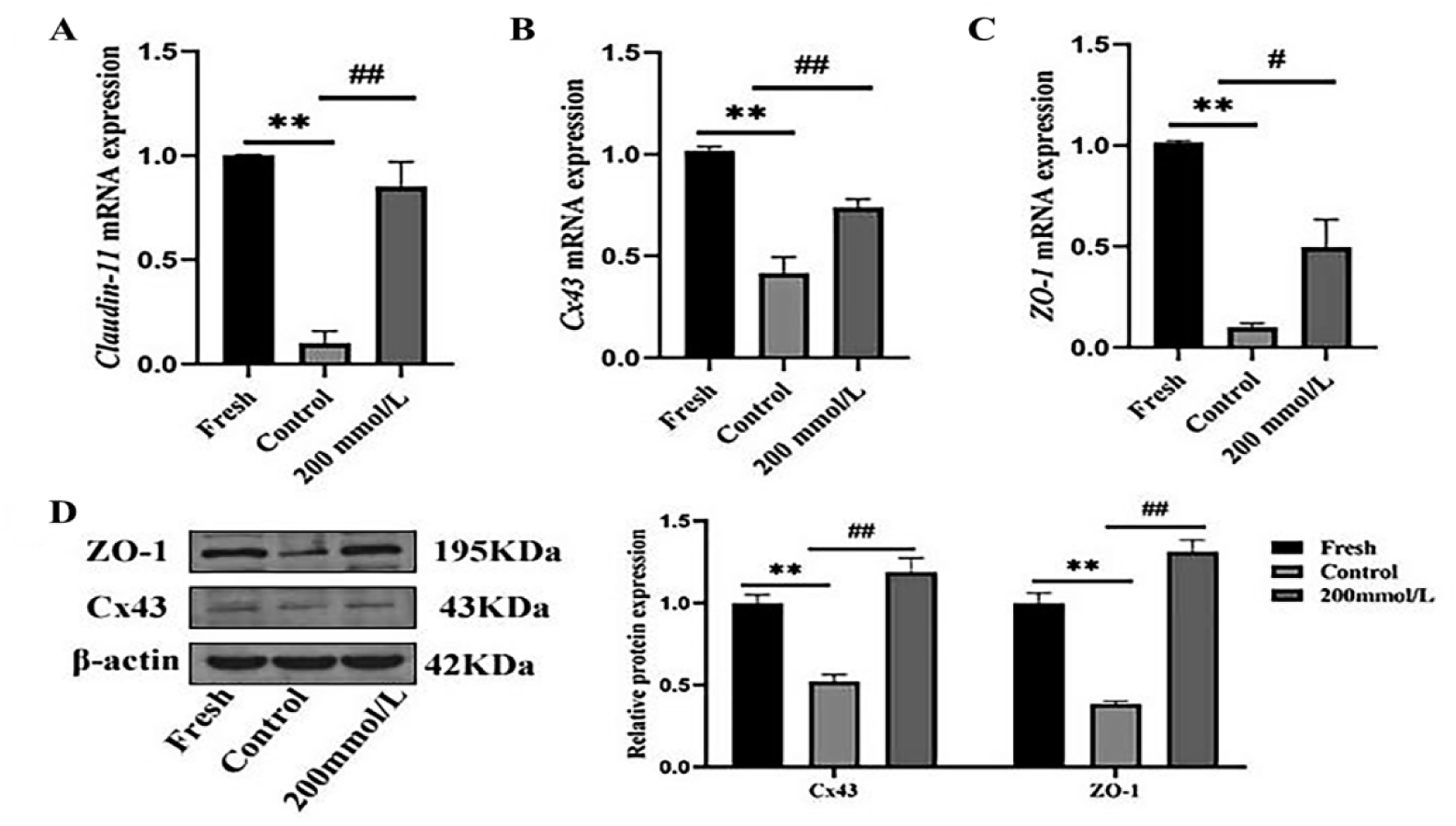
Effect of trehalose on BTB of frozen-thawed piglet testicular tissue. (A) *Claudin-11* mRNA expression; (B) *Cx43* mRNA expression; (C) *ZO-1* mRNA expression; (D) ZO-1 and Cx43 protein expression. The data are means ± SD (n = 3). \*\**P* < 0.01 indicates a statistically significant difference vs Fresh group; ^#^*P* < 0.05 and ^##^*P* < 0.01 indicates a statistically significant difference vs Control group.

### 3.7. The Effects of Trehalose on Spermatogenesisin of Frozen-thawed Piglet Testicular Tissue

Spermatogenesis is tightly coordinated by various cells, hormones, genes, and epigenetic factors, with crosstalk among signaling pathways playing a central role. Therefore, qRT-PCR and Western blot were used to detect the mRNA expression of *AKAP4*, *CFAP44*, *DDX25*, *DYP19L2*, and *HMGB2*, as well as the protein expression of AKAP4 and HMGB2, respectively. The results are shown in Figure 3-8. Compared with the Fresh group, mRNA expression of *AKAP4*, *CFAP44*, *DDX25*, *DYP19L2*, and *HMGB2* in frozen-thawed piglet testicular tissue was significantly reduced (*P* < 0.01); HMGB2 protein expression was significantly reduced (*P* < 0.01), and AKAP4 protein expression was significantly reduced (*P* < 0.05). Compared with the Control group, 200 mmol/L trehalose significantly increased the mRNA expression of *AKAP4*, *DDX25*, *DYP19L2*, and *HMGB2*, and the protein expression of AKAP4 and HMGB2.

**Figure 3-8.**
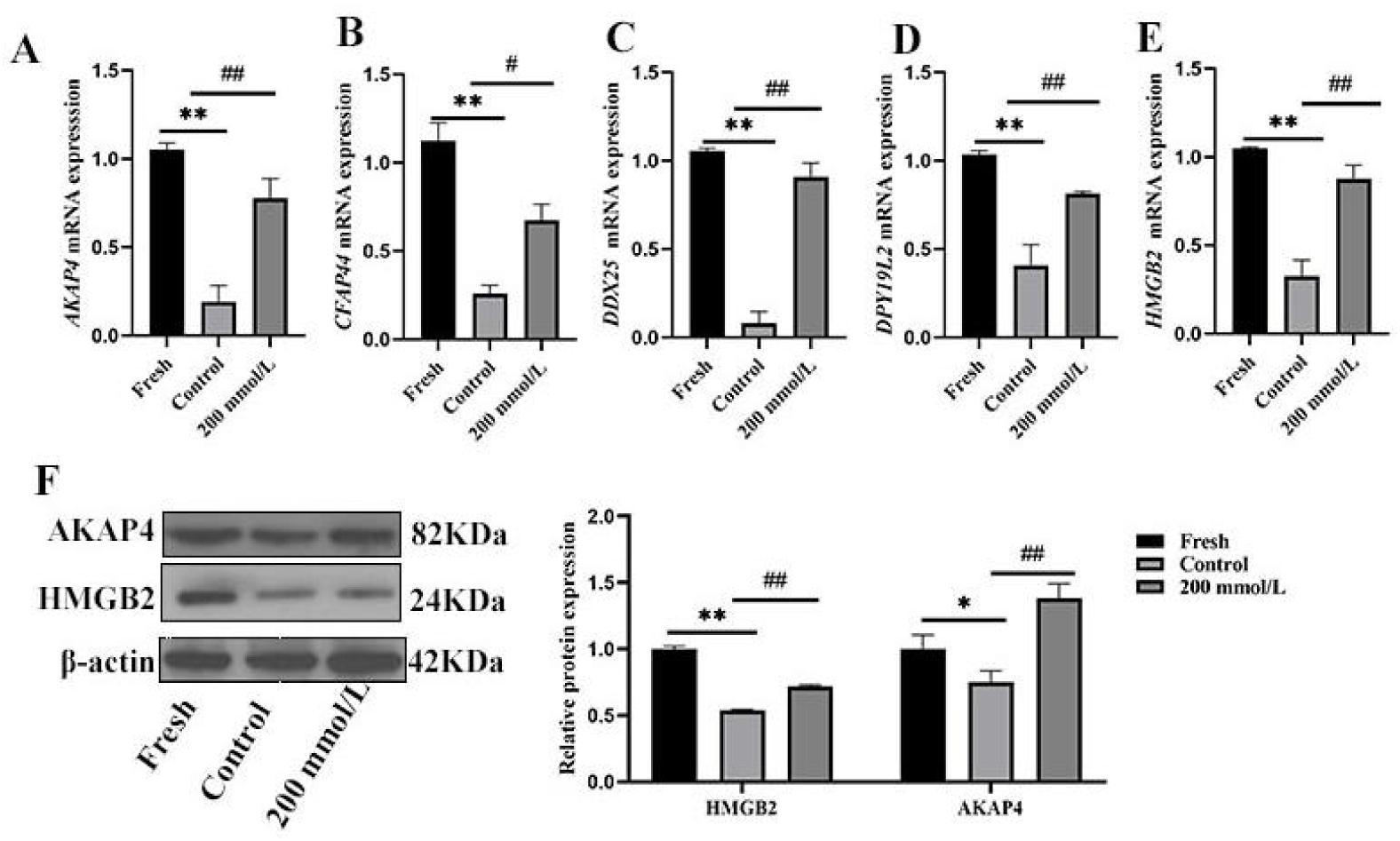
Effect of trehalose on spermatogenesis-related factors of frozen-thawed piglet testicular tissue. (A) *AKAP4* mRNA expression; (B) *CFAP44* mRNA expression; (C) *DDX25* mRNA expression; (D) *DPY19L2* mRNA expression; (E) *HMGB2* mRNA expression; (F) AKAP4 and HMGB2 protein expression. The data are means ± SD (n = 3). \**P* < 0.05 and \*\**P* < 0.01 indicates a statistically significant difference vs Fresh group; ^#^*P* < 0.05 and ^##^*P* < 0.01 indicates a statistically significant difference vs Control group.

### 3.8. DEGs Analysis

In the Fresh vs. Control group, 3,567 DEGs (1,777 upregulated, 1,790 downregulated) were found (Figure 3-9A); in the Control vs trehalose-treated group, 845 DEGs (624 upregulated, 221 downregulated) were found (Figure 3-9B). By comparing DEGs between the Fresh and Control groups, and between the Control and trehalose-treated group, the regulation of trehalose-responsive genes was analyzed. There were 89 downregulated genes in the Control group, but the genes upregulated in the trehalose-treated group. And 90 genes upregulated in the Control group, but the genes downregulated in the trehalose-treated group (Figure 3-9C).

**Figure 3-9.**
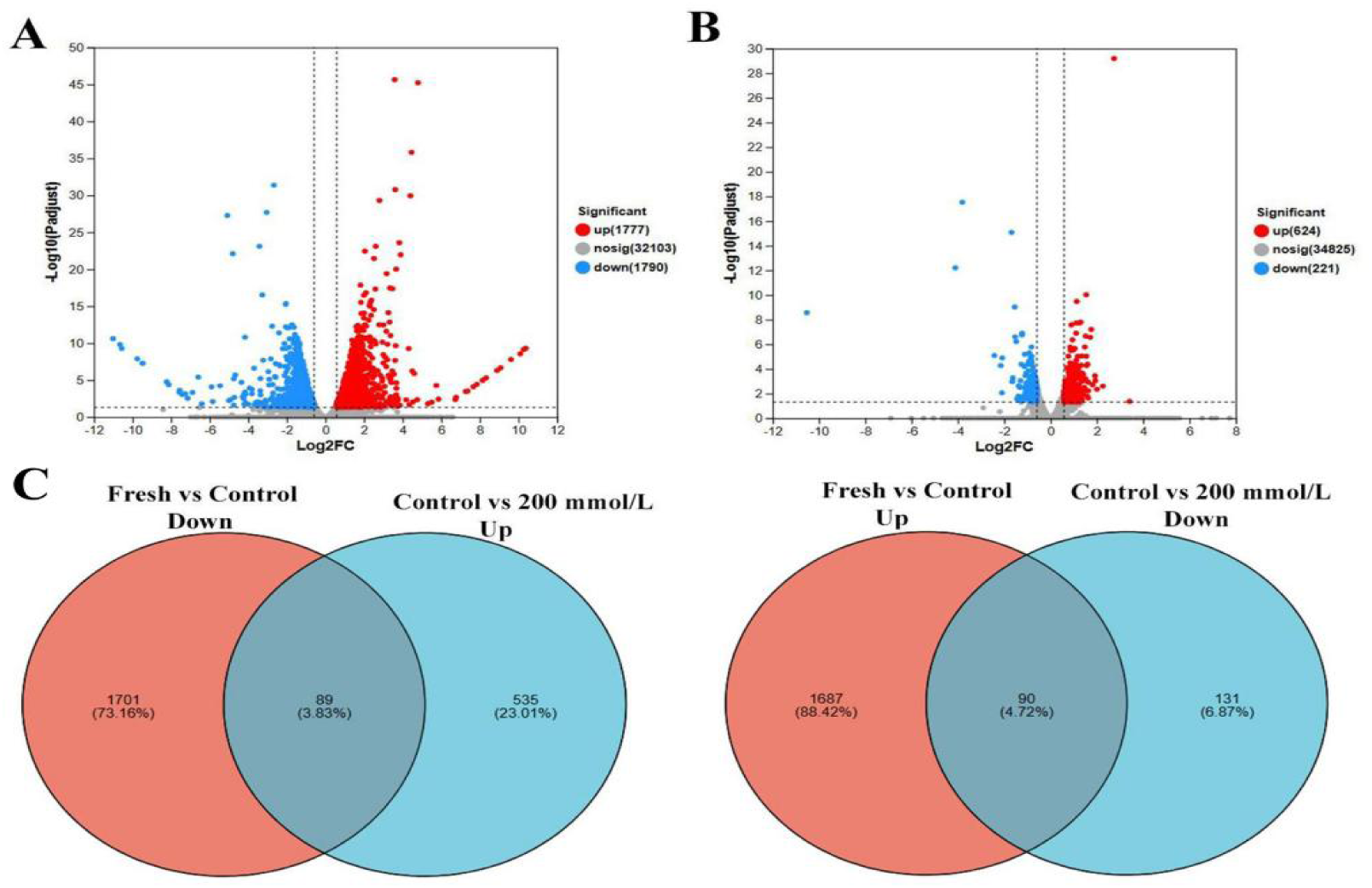
Expression difference volcano plot. (A) Fresh vs Control; (B) Control vs 200 mmol/L; (C) Venn graph of trehalose induced DEGs.

### 3.9. GO Enrichment Analysis of Trehalose-Regulated DEGs

GO comprises three categories: BP, CC and MF. GO enrichment analysis was performed on the downregulated DEGs, with statistical significance determined at *P* adjust < 0.05. In Figure 3-10, *P* < 0.01. After GO analysis it was revealed that trehalose-regulated DEGs were significantly enriched in mitochondrial-related BP, including respiratory electron transport chain (GO:0022904) and mitochondrial electron transport (GO:0006120). CC were predominantly enriched in the inner mitochondrial membrane protein complex (GO:0098800) and mitochondrial inner membrane (GO:0005743), among others. MF were primarily enriched in electron transport activity (GO:0009055) and NADH dehydrogenase activity (GO:0003954). Among these, *ND2*, *ND5*, *ND6*, and *CYTB* are all conserved gene encoded by the mitochondrial genome, while *COX2* is a key subunit of cytochrome c oxidase. The concurrent upregulation of these genes suggests that trehalose exerts its anti-apoptotic and protective effects by improving the dysfunction of cellular oxidative phosphorylation caused by frozen-thawed cycles through the synergistic regulation of the interaction between nuclear and mitochondrial genes.

**Figure 3-10.**
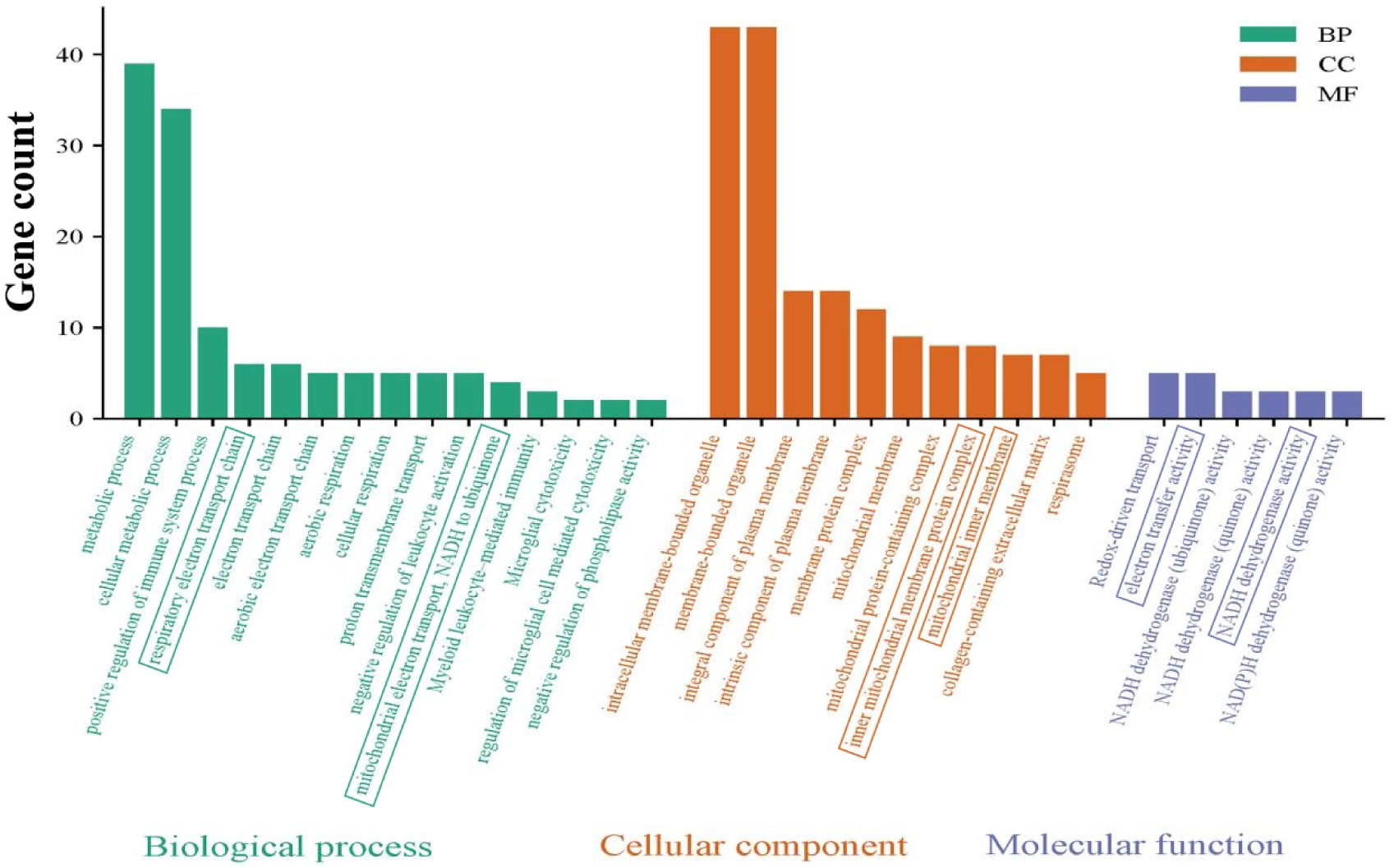
GO enrichment of trehalose-regulated DEGs.

### 3.10. KEGG Enrichment Analysis of DEGs Modulated by Trehalose

Oxidative phosphorylation (map00190) and Thermogenesis (map04714) were identified as the core pathways mediating trehalose function (Table S3). These pathways encompass key genes encoding subunits of the mitochondrial electron transport chain (mETC), including: *ND2*, *ND5*, *ND6* (Complex I); *CYTB* (Complex III); *COX2* (Complex IV); and *ATP6*, *ATP8* (Complex V). Frozen-thawed injury downregulated the expression of these ETC genes, and trehalose treatment significantly upregulated their expression (Table 3-1), indicating its role in potentially restoring ETC function, thereby safeguarding cellular energy supply and improving mitochondrial integrity. Concurrently, significant enrichment of multiple neurodegenerative disease pathways-such as Parkinson’s disease (map05012), Alzheimer’s disease (map05010), and Huntington’s disease (map05016), was revealed using KEGG analysis, highlighting an anti-apoptotic mechanism of trehalose (Table S3). A core pathology in these diseases is mitochondrial dysfunction, which was reversed by trehalose, suggesting that the molecular damage from freezing converges on mitochondrial dysfunction pathways shared with neurodegenerative diseases. Thus, trehalose likely protects mitochondria by modulating these common deleterious signaling cascades.

**Table 3-1.**
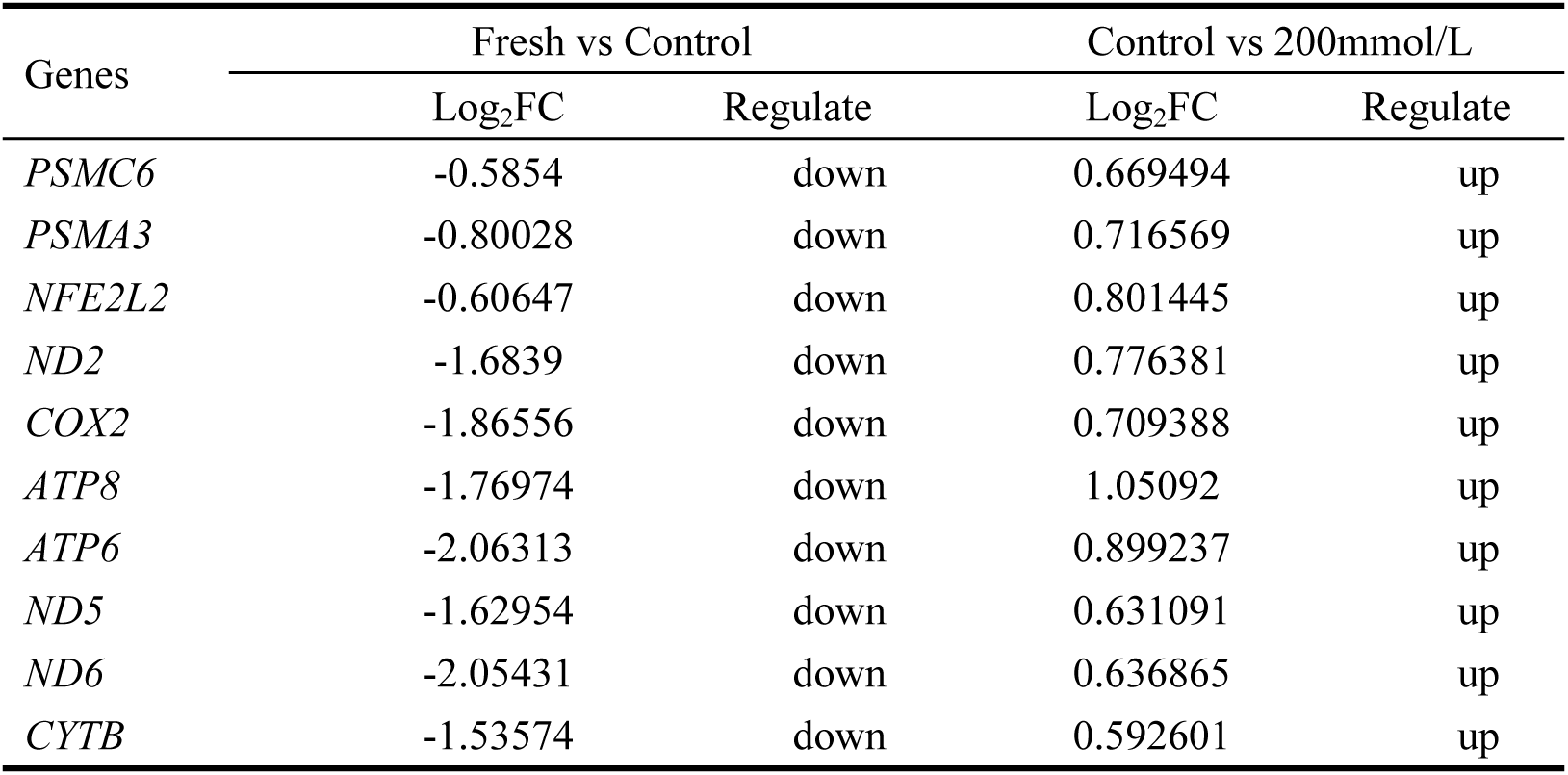
KEGG enrichment of trehalose-regulated DEGs.

### 3.11. qRT-PCR Confirmation of DEGs

GO and KEGG analyses revealed common functional categories, comprising proteasome function (*PSMC6*, *PSMA3*), antioxidative response (*NFE2L2*), and the mitochondrial respiratory chain/oxidative phosphorylation system (*ND2*, *COX2*, *ATP8*, *ATP6*, *ND5*, *ND6*, and *CYTB*) in Table 3-1. Building on this, RT-qPCR was used to validate the accuracy of the transcriptomic sequencing results at the molecular levels. The results show that, with the exception of *COX2*, all other variables showed statistically significant differences (Figure 3-11). High consistency with transcriptomic sequencing data was showed in the validation results, confirming the reliability of sequencing outcomes. These findings indicate that trehalose modulates mitochondrial function-related genes through multiple targets. Specifically, trehalose enhances mitochondrial antioxidative defense, while maintaining protein homeostasis primarily through mitochondrial-encoded respiratory chain subunits.

**Figure 3-11.**
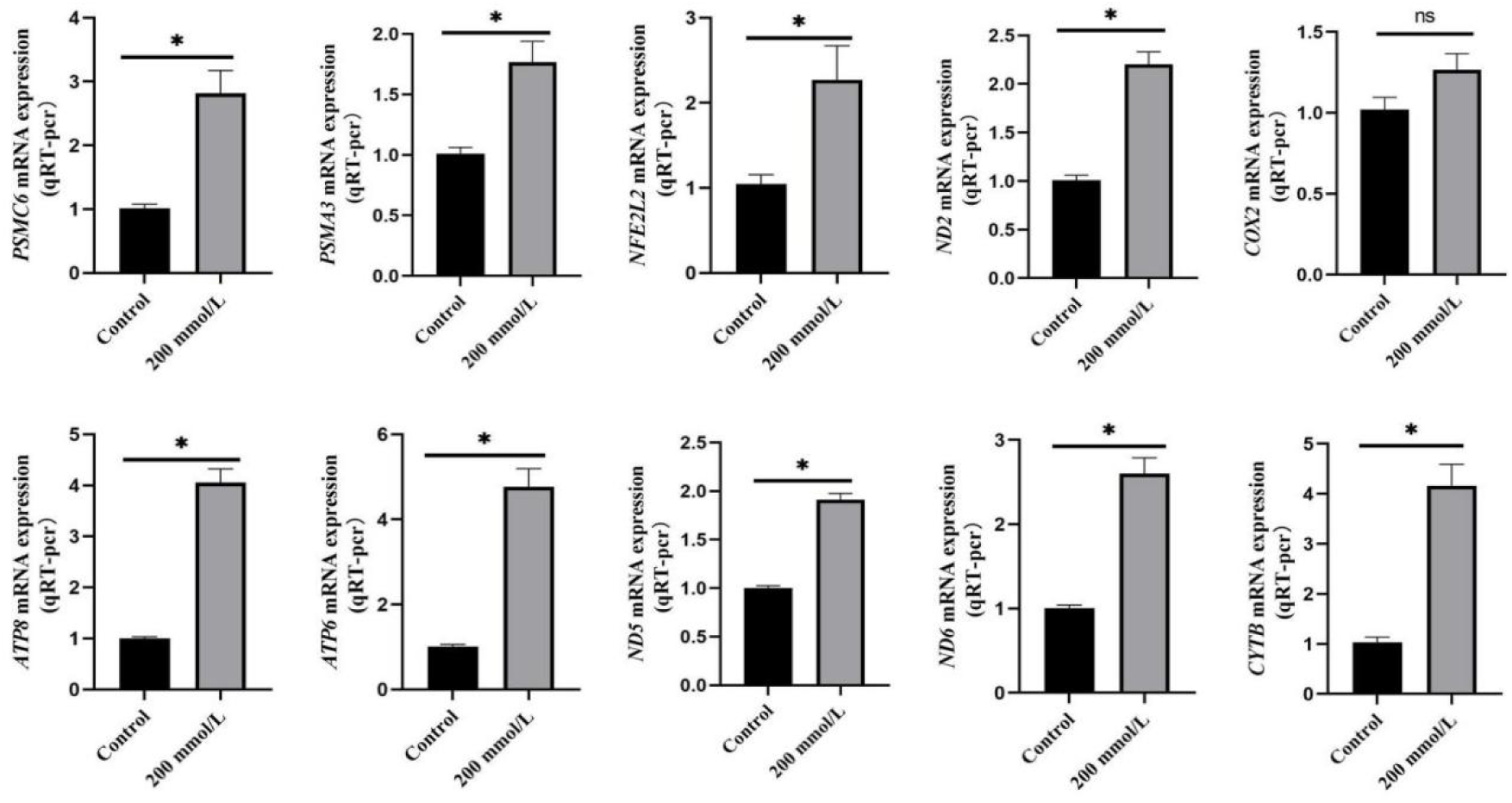
Validation of DEGs using qRT-PCR. The data are means ± SD (n = 3). \**P* < 0.05 indicates a statistically significant difference.

## 4. Discussion

We adopted slow freezing for porcine testicular tissue preservation based on prior comparative evaluations confirming its balanced operational feasibility and reduced cryoprotectant toxicity relative to vitrification [19–21], and supplemented the base DMSO-containing medium with trehalose, a widely validated non-permeable cryoprotectant that synergizes with permeable agents to mitigate ice crystal injury and osmotic stress [10]. Integrated analyses of post-thaw viability, apoptosis, histomorphology, and ROS levels consistently identified 200 mmol/L trehalose as the optimal concentration: this dose maintained cell viability and seminiferous tubule integrity comparable to fresh controls, suppressed apoptosis, and reduced oxidative stress, in line with prior reports on porcine spermatogonial stem cell cryopreservation [22]. Lower trehalose concentrations (≤ 100 mmol/L trehalose) failed to adequately limit ice nucleation, while higher concentrations (≥ 400 mmol/L trehalose) induced osmotic imbalance and cytotoxic ROS accumulation despite trehalose’s inherent antioxidant properties [23], ultimately exacerbating tissue damage. These protective effects are attributable to trehalose’s dual capacities to replace intracellular water to suppress ice formation and stabilize cell membranes via osmotic regulation [24], as well as enhance endogenous antioxidant defenses through Keap1-NFE2L2 pathway activation [25], supporting its utility as a safe, effective adjuvant for porcine testicular tissue cryopreservation.

Extensive studies demonstrate that freezing-induced ice crystal formation triggers apoptosis and pathophysiological changes, ultimately leading to reproductive dysfunction. Although trehalose exhibits beneficial biological functions, such as enhancing antioxidative enzyme activity and improving post-thaw tissue and cell quality, during the cryopreservation of various cell types and organisms, its effects on testosterone synthesis, laminin expression, and key genes involved in spermatogenesis within testicular tissue remain unexplored. As the testis is a critical site for androgen secretion, BTB establishment, and spermatogenesis, the effects of trehalose on the secretory function and expression of related key genes in frozen-thawed piglet testicular tissue was specifically investigated in this study. Previous studies have shown that freezing testicular tissue induces LCs apoptosis, resulting in reduced testosterone secretion [18,26]. Xi et al. [6] reported that enhancing LCs viability after cryopreservation of goat testicular tissue increases testosterone secretion. The results of the present study corroborated these findings: the frozen control group exhibited decreased testosterone secretion and downregulated mRNA expression levels of *CYP17A1*, *CYP11A1*, and *StAR*, indicating that freezing may inhibit testosterone synthesis by impairing the expression of key genes in the synthesis pathway. Trehalose restored testosterone secretion levels and the expression of these genes, suggesting that its reproductive protective effect is mediated, at least in part, by enhancing cell viability and reducing apoptosis. Beyond maintaining endocrine function, the structural integrity of the testis is highly dependent on the BTB. Located at the basement membrane of the seminiferous epithelium, the BTB is comprised of tight junctions, gap junctions, desmosomes, and focal adhesions [27]. Core tight junction proteins include claudins, occludin, and zonula occludens (ZO) proteins, wherein ZO-1 connects the actin cytoskeleton to transmembrane proteins such as claudin and occludin to maintain structural stability [28]. The claudin family is widely involved in the regulation of various tissues, and Claudin-11 is a key tight junction protein for the BTB; its knockout in mice leads to infertility [28–30]. Gap junctions are formed by members of the connexin (Cx) family, among which Cx43 is the most abundant [31]. In this study, the mRNA expression levels of *Claudin-11*, *Cx43*, and *ZO-1* were significantly downregulated in frozen-thawed piglet testicular tissues. In contrast, 200 mmol/L trehalose significantly upregulated the mRNA levels of *Cx43* and *ZO-1* compared to the frozen control group. These findings were subsequently validated using Western blot analysis. This suggests that trehalose acts as a membrane stabilizer, displacing intracellular water to prevent cell rupture caused by ice crystals, thereby playing a crucial role in maintaining the adhesive capacity of junctional proteins and the structural scaffold of the seminiferous tubules [24].

Spermatogenesis involves spermatogonial stem cell differentiation within seminiferous tubules, regulated by cells, hormones, genes, and epigenetic factors via crucial signaling crosstalk [32 – 33]. *DDX25*, a testis-specific DEAD-box family member expressed in meiotic and haploid germ cells, plays a key role in spermatogenesis [34]. *HMGB2*, an HMG family protein involved in transcription, replication, recombination, and repair, negatively regulates spermatid apoptosis [35]; Sugita et al. [36] Found that HMGB2 deficient mice exhibit reduced testicular weight, fewer seminiferous tubules, and increased apoptosis, with atrophic tubules containing only Sertoli cells (SCs). *DPY19L2*, a spermatid-specific transmembrane protein, is essential for sperm head elongation and acrosome formation [37]; its reduced expression in mice exposed to toxic environments inhibits acrosome biogenesis [38]. *CFAP44*, a flagellar protein, organizes the sperm flagellar axoneme, and its deficiency leads to male infertility with multiple flagellar abnormalities [39]. *AKAP4*, a major component of the sperm fibrous sheath, interacts with A-kinase anchoring protein 3 to mediate sperm phosphorylation modifications, regulating capacitation-related signaling and metabolism; both proteins are key structural determinants of flagellar sperm motility, and their absence results in flagellar anomalies and infertility [40,41]. Consistent with prior evidence that cryopreservation reduces spermatogenesis-related gene expression [42], our results show that frozen-thawed tissues exhibited significantly downregulated mRNA levels of *AKAP4*, *CFAP44*, *DDX25*, *DPY19L2*, and *HMGB2*, alongside reduced protein expression of HMGB2 and AKAP4. This indicates that cryopreservation alters genes pivotal to spermatogenesis. Notably, trehalose supplementation significantly increased the expression of these genes compared to the frozen control group, demonstrating its spermatogenic protective effect during testicular tissue cryopreservation.

DEGs in piglet testicular tissue following trehalose supplementation were analyzed to elucidate the molecular mechanism underlying these phenomena. Transcriptomic profiling revealed significant alterations in mitochondrial dynamics, as freezing disrupted the homeostatic balance between mitochondrial fusion and fission—a finding corroborated by altered expression of key respiratory chain components such as Complex I subunits *ND5*/*ND6* and *COX2* [43]. Concurrently, KEGG enrichment analysis showed that downregulated DEGs were predominantly associated with oxidative phosphorylation, ROS metabolism, and neurodegenerative disease-related pathways, all of which are closely linked to mitochondrial dysfunction and redox imbalance. This enrichment pattern closely resembled the findings of Wang et al. [44] on fish sperm cryopreservation, suggesting a conserved mechanism of cryo-injury across species. At the molecular level, the significant downregulation of the 26S proteasome subunits *PSMC6* and *PSMA3* in the frozen control group is particularly noteworthy. Given that the 20S core proteasome (one of the two 26S subcomplexes) is essential for meiotic progression and the 26S proteasome is involved in sperm capacitation, their suppression indicates disrupted protein quality control and cell cycle regulation [45,46]. In contrast, trehalose significantly upregulated the core antioxidative transcription factor *NFE2L2*, indicating robust activation of the endogenous antioxidative defense system [47,48]. This upregulation enhances glutathione synthase expression and maintains mitochondrial glutathione stability, playing a pivotal role in mitochondrial redox homeostasis [49]. Furthermore, trehalose preserved the normal expression of electron transport chain components such as *CYTB*, *ATP6*, and *ATP8*, thereby stabilizing mitochondrial membrane potential and ensuring the structural and functional integrity of the mitochondrial reticulum, which ultimately augmented the tissue’s capacity to resist cryo-induced oxidative stress [50,51].

Based on the aforementioned molecular mechanism investigations, the authors posit that mitochondrial dysfunction serves as the central hub connecting freezing-induced physical stress to subsequent physiological decline. Cimolai et al. [52] demonstrated that the ROS burst and excessive accumulation within the mitochondrial respiratory chain, surpassing the scavenging capacity of the antioxidative system, may lead to mitochondrial dysfunction, including impaired respiratory complex activity and reduced ATP levels. Another study indicated that mitochondria are closely associated with steroid biosynthesis, and that disruption of steroid biosynthesis induced by mitochondrial dysfunction may lead to male reproductive dysfunction [53]. 200 mmol/L trehalose restored this balance, not only preserving the integrity of the outer mitochondrial membrane but also potentially alleviating damage through the induction of mitophagy [54,55]. This constitutes the core mechanism underlying its cryoprotective effect. As Tiwari et al. [56] emphasized, targeted antioxidative strategies are crucial for maintaining sperm quality. This study expands upon this theory by demonstrating that trehalose stabilizes the homeostasis of mitochondrial fusion and fission, promoting cell proliferation and cell cycle progression, thereby regulating DNA transcription and gene expression.

In summary, this study indicates that trehalose primarily exerts its cryoprotective effect by stabilizing mitochondrial dynamics. While previous studies focused solely on mitochondrial damage in spermatozoa [57], this work highlights the importance of mitochondrial homeostasis within the broader context of testicular tissue architecture—encompassing somatic cell (Leydig) function through to germ cell spermatogenesis. By preserving the balance of mitochondrial fusion and fission, trehalose not only enhances ATP production but also strengthens the antioxidative defense system. Future research should strive to translate these molecular findings into clinical outcomes, ensuring the restoration of reproductive function after transplantation.

## Supplementary Materials

The supporting information can be downloaded at:

## Author Contributions

H.Y. and L.N. were responsible for experimental design, sample, data analysis, and writing of the manuscript. L.J. and W.Y were responsible for sample collection and data analysis. W.X and L.X were responsible for the discussion about the experimental design. And all authors contributed to the article and approved the submitted version. X.C, Z.J and H.C reviewed and edited the manuscript.

## Funding

This research was funded by Nanning City Scientific Research and Technology Development Plan (20253037).

## Institutional Review Board Statement

This study strictly adheres to the relevant regulations and guidelines of the Guangxi University Laboratory Animal Ethics Committee for animal husbandry and experimentation. Animal ethics review number: Gxu2022-211.

## Data Availability Statement

The original contributions presented in the study are included in the article.

## Acknowledgments

The authors acknowledge Guangxi Yibiantianyuan Breeding Stock Co., Ltd. for supplying the biological materials (piglet testes) required for this study.

## Conflicts of Interest

“The funders had no role in the design of the study; in the collection, analyses, or interpretation of data; in the writing of the manuscript; or in the decision to publish the results.”

## Notes

### Competing Interest Statement

The authors have declared no competing interest.

